# Cross-pollination is more important than gibberellin application in the germination of the everlasting flowers of *Janeirona carrasqueira* (morphotype of *Comanthera bisulcata*)

**DOI:** 10.1101/2023.08.03.551682

**Authors:** Maria Luiza de Azevedo, Maria Neudes Sousa de Oliveira, André Rodrigo Rech, Jose Carlos Barbosa, Eric Bastos Gorgens, Evandro Luiz Mendonça Machado, Israel Marinho Pereira

## Abstract

The “sempre-vivas” are plants known as “everlasting flowers” because they have inflorescences that retain the appearance of living structures even after being harvested and dried. Given their ornamental potential, some species are widely exploited commercially, and their production occurs almost exclusively through extractivism. However, data on their pollination, reproduction, and germination are limited. Extractivism is an activity of great importance for the residents of the Espinhaço Meridional municipalities, and research is essential for establishing plans and proposals for the management of these native species. Given this scenario, we studied the pollinators and the importance of cross- and autogamy for reproductive success of *Janeirona carrasqueira* (morphotype of *Comanthera bisulcata* (Körn) L.R. Parra & Giul), commercially collected in Diamantina, Minas Gerais. We also evaluated germination and the influence of the exogenous application of gibberellin (GA3) on seeds. Pollinators were observed during the flowering period in beds cultivated with this species in the native Campo Rupestre area of Diamantina, Minas Gerais. The inflorescences of the species were collected from two different environments: one isolated from pollinating insects (inside the greenhouse), and the other open in the presence of pollinators. The seeds were removed from the flower heads and subjected to exogenous application of gibberellin (0–control and 500 ppm). The flowers showed a generalist pollination strategy, including visitors who collected pollen and nectar as well as those who exclusively collected nectar. For germination, the most influential factor was cross-pollination and not gibberellin application. We conclude that the pollination system represents a determining mechanism and is a very important factor to be considered in management plans aimed at species conservation.

**Highlights:**

– *Comanthera bisulcata* employs a generalist pollination strategy.
– The pollination system is a crucial mechanism for germination.
– Cross-pollination is more important than gibberellin application
– Reproduction is a crucial aspect to be taken into account in the management plans of the species.

## 1. Introduction

The Brazilian Campo Rupestre is among the most diverse environments on the planet, and the Diamantina Plateau region, where this phytophysiognomy predominates, has recently been considered an exclusive province of biodiversity (Colli-Silva et al., 2019). This environmental peculiarity is due to the high diversity of plants (∼15% of the Brazilian flora in less than 1% of the Brazilian territory) and the high levels of endemic species (Silveira et al., 2016; Fernandes et al., 2020; Vasconcelos et al., 2020). Among many drivers of this biological diversity is the mosaic of soil types that results in different nutrient acquisition strategies by vegetation (Castro et al., 2018; Abrahão et al., 2019). Some species of the family Eriocaulaceae are among the emblematic plants growing in this diversity of soils, especially in those that are very poor in nutrients (Costa et al., 2008; Giulietti et al., 2012). Although poorly studied, the pollination and reproductive systems in the family seems also to be very diversified (Ramos et al., 2005; Del-Claro et al., 2019; Andrino et al., 2022; Martins-Junior et al. 2022).

The Eriocaulaceae family has approximately 1100 species distributed across 10 genera with a pantropical distribution, although the Serra do Espinhaço region homes the vast majority of existing species (Giulietti et al., 2012). Its representatives are characterized by the presence of capitulum inflorescence (Stützel and Trovó, 2013). Many of these plants are known as ‘sempre-vivas’ (everlasting flowers) because their inflorescences retain the original shape for a long period after being collected and dried (Parra et al., 2010). Around 30 species in the family are already endangered (MMA, 2022) and there is even record of extinction in nature in Japan (Tanaka et al., 2015), making the studies on seed production and germination urgently demanded for Eriocaulaceae.

Eriocaulaceae members belonging mainly to *Syngonanthus* and *Comanthera* genera, are economically important ornamental plants (Parra et al., 2010). In many communities in Serra do Espinhaço, flower collection is an important source of income for families (Oliveira et al., 2014). The practice of collection, traditionally known as “flower picking” by extractive communities, has recently been recognized by the FAO-UN as the first existing world agricultural heritage in Brazil (FAO, 2020; Borges and Branford, 2021).

Owing to their significant economic and social impact (Gonçalves-Magalhães et al., 2021), combined with the endemic distribution of many species with everlasting flowers, measures to ensure their preservation are necessary. These measures include studying the reproductive conditions (pollination, crossing, and germination systems) of commercialized species to improve *ex situ* cultivation techniques, and design adequate management strategies for native populations subjected to extractivism (Schmidt et al., 2008; Horiuchi et al., 2020). Currently, the lack of knowledge about reproductive systems is considered a major problem in implementing appropriate conservation measures (Horiuchi et al., 2021).

Only few studies have been conducted on the reproductive and germination characteristics of commercially available everlasting flower species. The huge heterogeneity in the germination rates of these species (ranging from 19% to 90%), as reported by previous studies, indicates the difficulty in achieving satisfactory germination rates for some species. (Ramos et al., 2005; Simões et al., 2007; Oriani et al., 2009; Oliveira and Garcia, 2011). Germination is a complex physiological process, affected by environmental factors and regulated by several hormones (Garcia et al., 2020). Among the hormones known to play a role in germination are gibberellins (Barreto et al., 2020), which are widely used to promote planting when germination is insufficient. The mechanism of action of gibberellins involves the induction of hydrolytic enzymes that facilitate the breakdown of the seed coat, mobilize storage reserves, and promote embryo growth (Sun et al., 2004; Pawłowski, 2009).The relative contribution of ecological (pollination) and physiological (hormones) factors to the process of germination of everlasting flowers is poorly understood. This information is vital for establishing strategies that assist in the conservation of these species.

To evaluate the effect of ecological factors (pollination) and physiological factors on the germination of everlasting flowers, we studied the species known as “*Janeirona carrasqueira”* (morphotype of *Comanthera bisulcata* (Körn) L.R. Parra & Giul). We sought to understand and identify the dependence of this everlasting flower on pollinators. We also studied the role of exogenous gibberellin (GA3), and the reproductive system in the germination and early development of seedlings.

## 2. Material and Methods

### 2.1. Characterization of the study area

The study was conducted at the Federal University of Vales do Jequitinhonha e Mucuri (UFVJM), Campus JK (18° 12′ 3″ S, 43° 34′ 31″ W, average altitude: 1,296 m). According to the Köppen classification, the climate in the region is Cwb, humid, temperate, with rainy summers and dry winters (Alvares et al., 2013). The average annual temperature is 18.3 °C, with an average minimum of 14.1 °C and an average maximum of 23.7 °C (INMET, 2020). The average annual precipitation is 1400 mm, and the rainy season occurs from October to March, representing 88% of the total annual precipitation (Vieira et al., 2010).

The university campus has an area of approximately 60 ha of native Rupestrian fields. The area is adjacent to Biribiri State Park, with preserved area of more than 16,000 ha housing the greatest diversity of everlasting flower species in the region (Echternacht et al., 2012; Andrino et al., 2015). In this study, plants up to 6 yrs old, grown in open-air managed beds, and plants grown in vases of 15 cm diameter, containing the same soil as the substrate were used. The plants in the vases were kept in a greenhouse closed to the side with a shade screen and a PVC plastic roof. The plants in the pots were the transplanted seedlings from the beds already established at the time of the experiment and also from first/second flowering. The beds were close to the greenhouse and were therefore subjected to the same microclimatic characteristics, except those influenced by the greenhouse itself.

### 2.2. The evaluated species

The “*Janeirona carrasqueira*” is also known as “daisy” (similar to the Asteraceae) and belongs to the genus *Comanthera*, and subgenus *Comanthera*. Two *C. bisulcata* morphotypes recognized are popularly referred to as “sempre-viva chapadeira” and “sempre-viva janeirona carrasqueira”. Both are morphologically characterized by their rosette-shaped growth habits, from which capitula-type inflorescences arise. The “*Janeirona carrasqueira*,” is among the commercialized everlasting flowers, and often collected from Diamantina-MG region between the months of January and February. It is considered a “quick flower,” a term popularly used to refer to capitula that remain in a marketable pattern for a short period, in relation to those of other commercialized species of the same genus.

### 2.3. Floral visitors and pollinators

Floral visitors were observed for three days during flowering in December 2019. The observation interval was 8 h (8:00 am to 4:00 pm), divided into 30 min blocks. The initial 10 min were used to count visitors, and the remaining 20 min were used to describe their behavior. Information on the period and duration of visits and the number of individuals/inflorescences visited at different times throughout the planted bed were recorded. Digital footage was captured to better visualize and understand the visitor behavior.

The distinction between pollinators and robbers was determined based on the foraging behavior displayed by visitors during the collection of floral resources and the nature of their contact with anthers and stigmas. In addition, we examined the presence of pollen adhered to the body to exclude floral visitors that could not be “*Janeirona carrasqueira*” pollen vectors. Based on the classification of visitor behavior, we divided them into nectar collectors, pollen collectors, or both. We then modeled the visit probability of these behaviors as a function of the time of day and number of flowers visited using a Generalized Linear Model (binomial GLM).

We also evaluated the location of adhesion and the load of pollen that adhered to the body surface. The pollen load adhered to the body surface is considered an indicator of the floral fidelity of visitors. For this purpose, the visitors collected in individual flasks were euthanized with fumes of ethyl acetate and the pollen on the body was removed with small cubes of glycerinated gelatin. Because their pollen morphology was known, *’Janeirona carrasqueira*’, pollen grains were distinguished from pollen grains from other plant species, and morphotypically quantified.

### 2.4. Collection of Inflorescences

Inflorescences were collected on the UFVJM campus at the two locations described above (bed and greenhouse). At Site 1, which was in an open environment, there were no barriers preventing floral visitors from accessing the inflorescences of the species (natural pollination treatment). At Site 2 (inside the greenhouse), the plants had no contact with floral visitors; therefore, the seeds produced were considered to originate from autogamy.

Floral chapters were collected in the first week of February 2020, when the scapes of different plants were randomly collected. In the Plant Physiology Laboratory, located in the Department of Agronomy at UFVJM, seeds were extracted from flower heads after approximately six months of storage at room temperature. Seeds from each treatment group were extracted by gently rubbing the floral heads with tweezers in a Petri dish. Subsequently, they were separated under a stereoscopic magnifying glass STEMI 2000-C/ (Zeiss, Germany), and malformed seeds were eliminated.

### 2.5. Germination Test

Seeds were subjected to asepsis in commercial sodium hypochlorite solution at a concentration of 2.5% (v/v) for 15 min and then rinsed twice in distilled water. Subsequently, the seeds were immersed in a solution of gibberellin (GA3) dissolved in distilled water at a concentration 500 ppm for 24 h at room temperature for imbibition to break dormancy. Seeds immersed plain distilled water served as the control. The seeds were then placed in Petri dishes containing two sheets of filter paper. They were then moistened daily with distilled water. The plates were incubated in a Mangelsdorf germinator, at 25±2°C, without artificial light supplementation. To reduce contamination, the filter paper and Petri dishes were disinfected with 70% alcohol.

The experimental design was completely randomized in a 2 × 2 factorial scheme (two collection environments in the presence or absence of GA3). Five replicates of 30 seeds each were used in each treatment. The evaluations were performed every five days under a stereoscopic magnifying glass. We evaluated the germination percentage, relative frequency of germination (Labouriau and Valadares, 1976) and postseminal development. We considered germinated seeds as those that presented a protrusion of the embryonic axis. The experiment was conducted for 52 d, till the beginning of seedling senescence was observed.

Statistical analyses were performed using R software version 4.2.2 (R Core Team, 2013). The effects of treatments on seed germination were analyzed using generalized linear mixed models (GLMM) implemented with the ‘glmer’ function of the R package ‘lme4’. For every seed (germinated or not), a binomial distribution was considered, and replication was treated as a random factor. Models were built considering different combinations: pollination * giberelin, pollination + giberellin, only pollination, only giberelin and a null model using only the intercept. We performed a Model Selection Analysis, taking into account the Akaike Information Criteria.

## 3. Results

### 3.1. Floral visitors and pollinators

We observed that ‘*Janeirona carrasqueira’* in Diamantina-MG presents synchronized flowering at the population level, but sequential at the individual level. This sequence is protandrous; that is, the male flowers open before the female flowers. Female flowers provide nectar as the only resource for visitors.

Insects of the Hymenoptera order were the ones that showed the highest frequency of visits to the *’Janeirona carrasqueira’* flowers (56.22 %), followed by Diptera (26.02%), Coleoptera (14.5%) and Lepidoptera (3.26%), showing great diversity in behavior. Visits by Lepidoptera were surprising, as the floral morphology of Eriocaulaceae generally does not favour insects with elongated mouthparts, such as moths and butterflies.

Most insects that visited flowers were small relative to the size of the inflorescence (Fig. 1), with mouthparts of different sizes and shapes. Visitors of the order Hymenoptera collected pollen and/or nectar, with quick visitation lasted between 3 to 6 s, proceeding sequentially to the next flower in the same flowerbed. Among these, Halictidae bees collected pollen exclusively during visits. Two different behaviors were recorded for the flies. In the first group, pollen was collected by Syrphidae flies. In the second group, other flies exclusively collecting nectar were recorded. Representatives of the order Coleoptera were recorded feeding on the pollen. *Pyrausta sp*. (Crambidae) was the only Lepidoptera recorded to feed on floral nectar from the flowers of the *’Janeirona carrasqueira*’.

**Figure 1.**
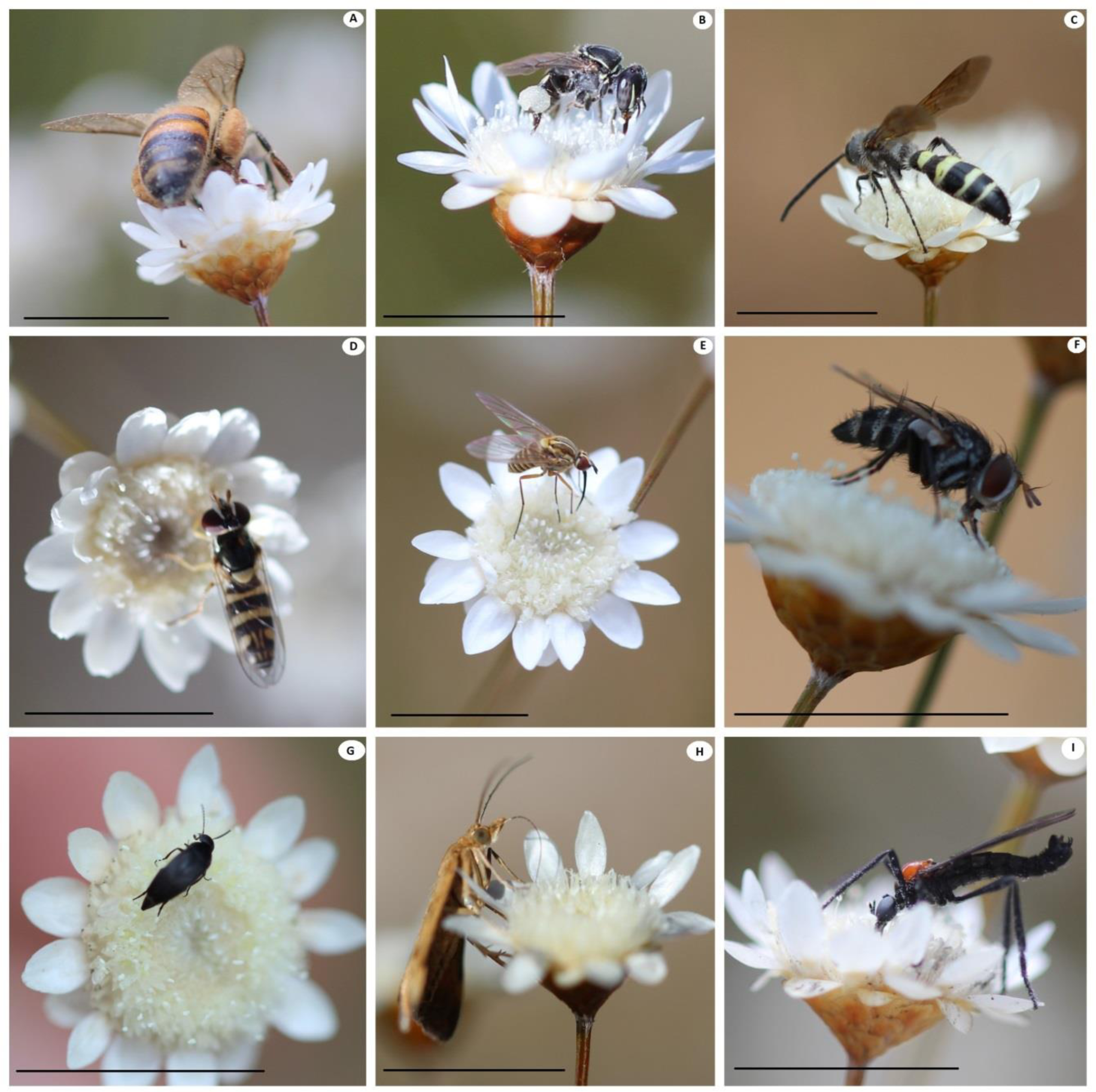
Sample of floral visitors of *Janeirona carrasqueira* (morphotype of *Comanthera bisulcata*) in Diamantina - MG. Bees: *Apis mellifera* (A); *Plebeia* sp. with polen (B); wasp (Vespidae) (C); Fly eating pollen (Syrphidae: Diptera) (D); Nectar-feeding mosquito (Diptera) (E); Fly eating nectar (Muscidae:Diptera) (F); Beetle (Coleoptera) (G); *Pyrausta* sp. (Crambidae) (H); *Plecia* sp. (Diptera) (I). Bars = 1 cm

The probability of visits for pollen collection increased by 25% between 1:00 pm and 2:00 pm and by 50% in the late afternoon; however, it was also related to the number of flowers visited (Interaction between time and number of visited flowers: χ²=5.82; p=0.015; Fig. 2A, 2B). Pollen collectors, essentially Apidae, carry out sequential visits, tasting different flowers (more than 12) on each flight to the flowerbeds. The highest richness of visitors (9 species) was observed in the morning when newly opened flowers provided nectar. Five floral visitors were recorded in the afternoon. The average abundances recorded in the 10 min intervals of observation in the morning and afternoon periods were 78 and 100 individuals, respectively. Although nectar and pollen were available in both periods, visitors such as Coleoptera were able to feed only on pollen in the afternoon. Nectar was explored by a greater number of species (richness), whereas pollen was used by a greater abundance of visiting insects.

**Figure 2.**
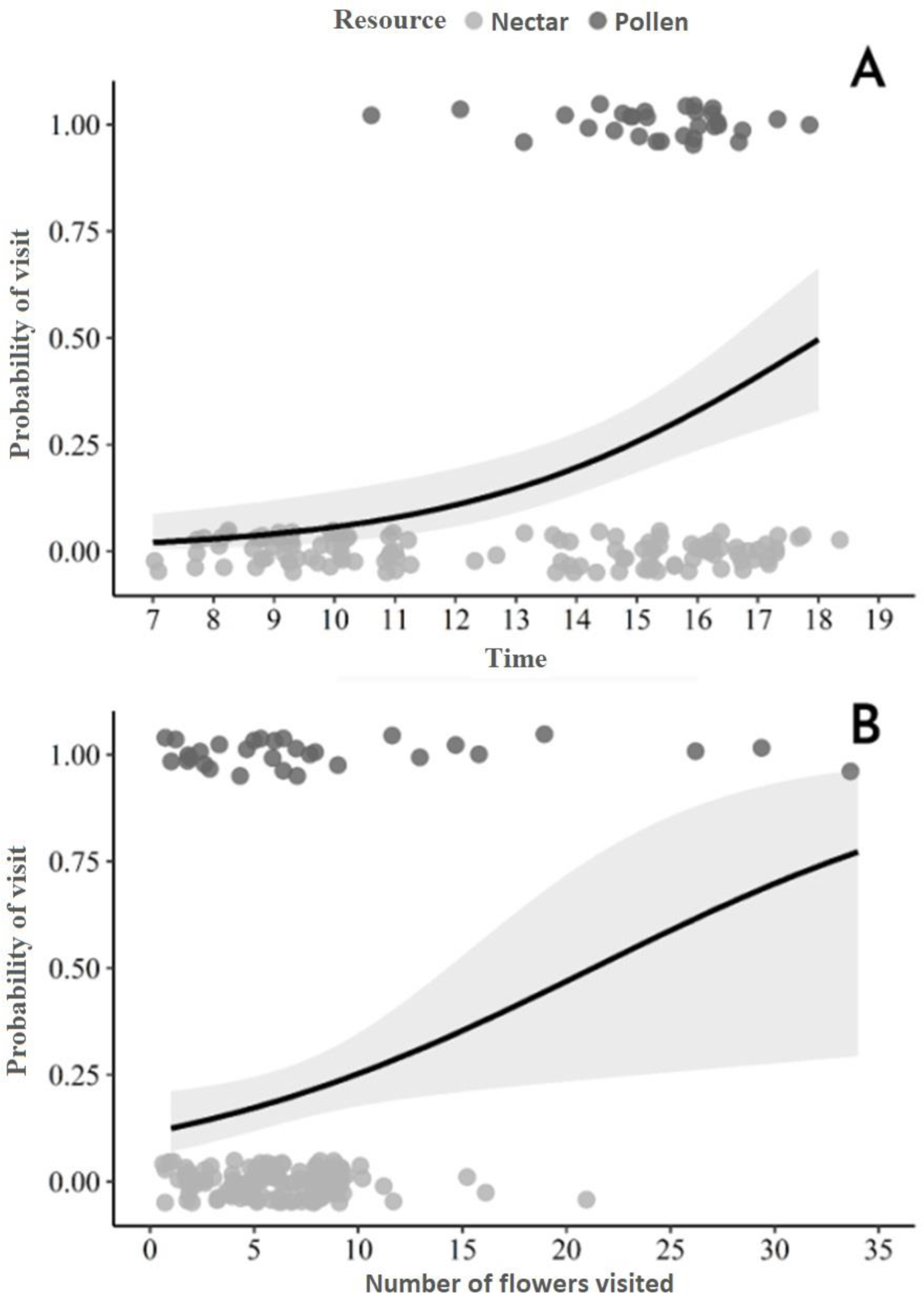
Generalized linear models of the probability of visits with collection of nectar and pollen resources as a function of visiting hours (A), and number of flowers visited (B).

A total of 70 floral visitors were collected, with an average of 23 pollen grains per visitor (SD = 36.07), ranging from 1 to 126 grains per animal. Among the species of flower visitors, 25 did not carry pollen from *’Janeirona carrasqueira’*. In addition to pollen from *’Janeirona carrasqueira*,’ pollen grains from over 18 species of plants were found in the floral visitors collected.

### 3.2. Germination test

The seeds of this species are small (millimeter-scale) and reddish-brown in color (Fig. 3A). Germination, characterized by protrusion of the embryonic axis (Fig. 3B) began seven days after the sowing of the collected seeds in open-air beds and in the greenhouse (Fig. 4). Protocorm differentiation to the polarized state (differentiation between leaf and root primordia) (Fig. 3C) was observed from day 12 onwards (Fig. 4). Throughout the germination process, leaves develop first, followed by the roots. The first (Fig. 3D) and second leaves (Fig. 3E) emerged after 17 and 27 d, respectively (Fig. 4). The maximum number of leaves emerged was three (Fig. 3F), which was observed after 42 days (Fig. 4). At 52 d, the beginning of senescence was observed in some seedlings, indicating depletion of their reserves.

**Figure 3.**
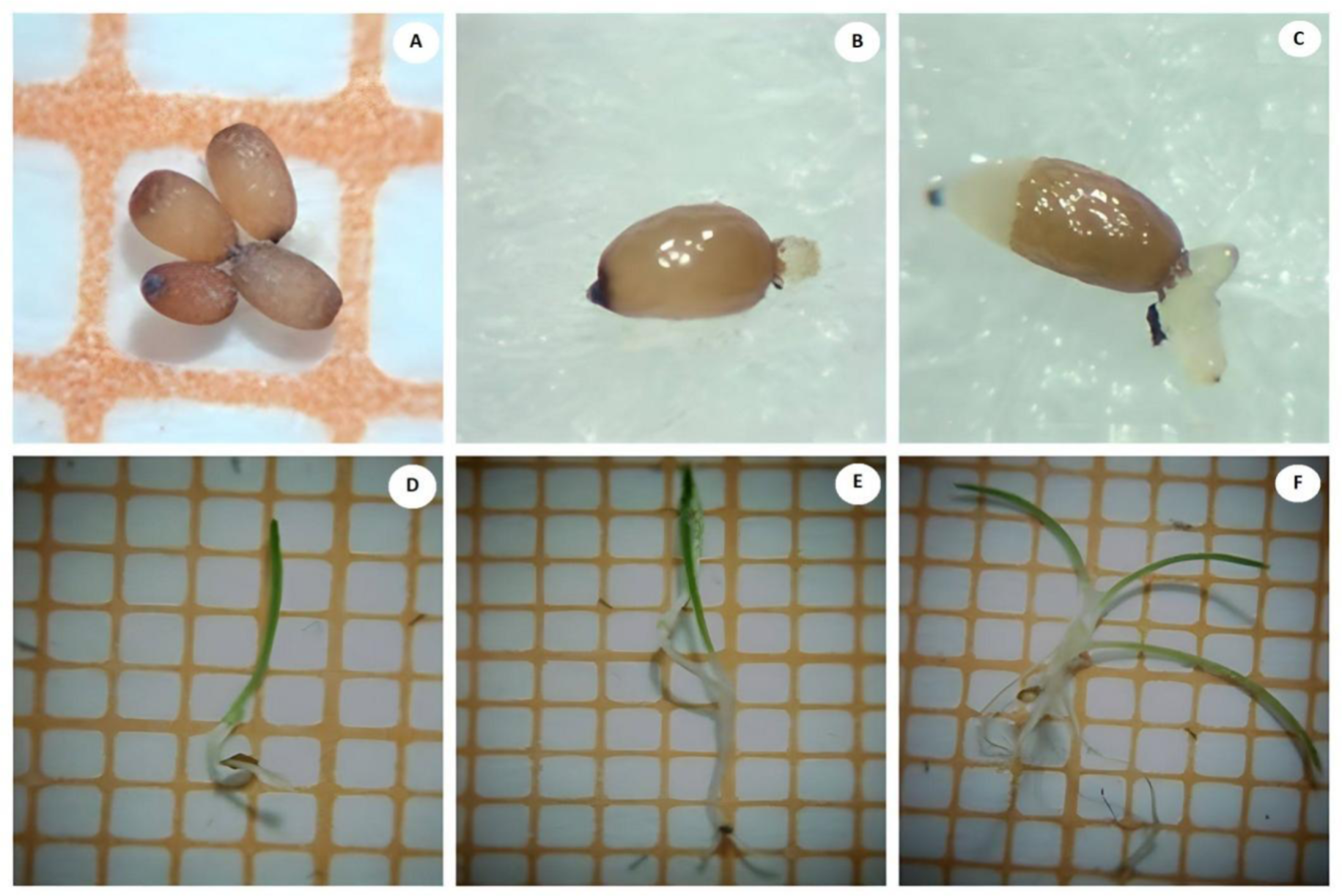
Seeds and seedlings of the “sempre-viva” *Janeirona carrasqueira* (morphotype of *Comanthera bisulcata*): (A) seeds collected in February; (B) protrusion of the embryonic axis, at 7 d after sowing; (C) polarization at 12 d after sowing, (D) 1^st^ leaf, 17 d after sowing; (E) 2^nd^ leaf, 27 dafter sowing; (F) 3^rd^ leaf, 47 d after sowing. Background squares = 1mm^2^.

**Figure 4.**
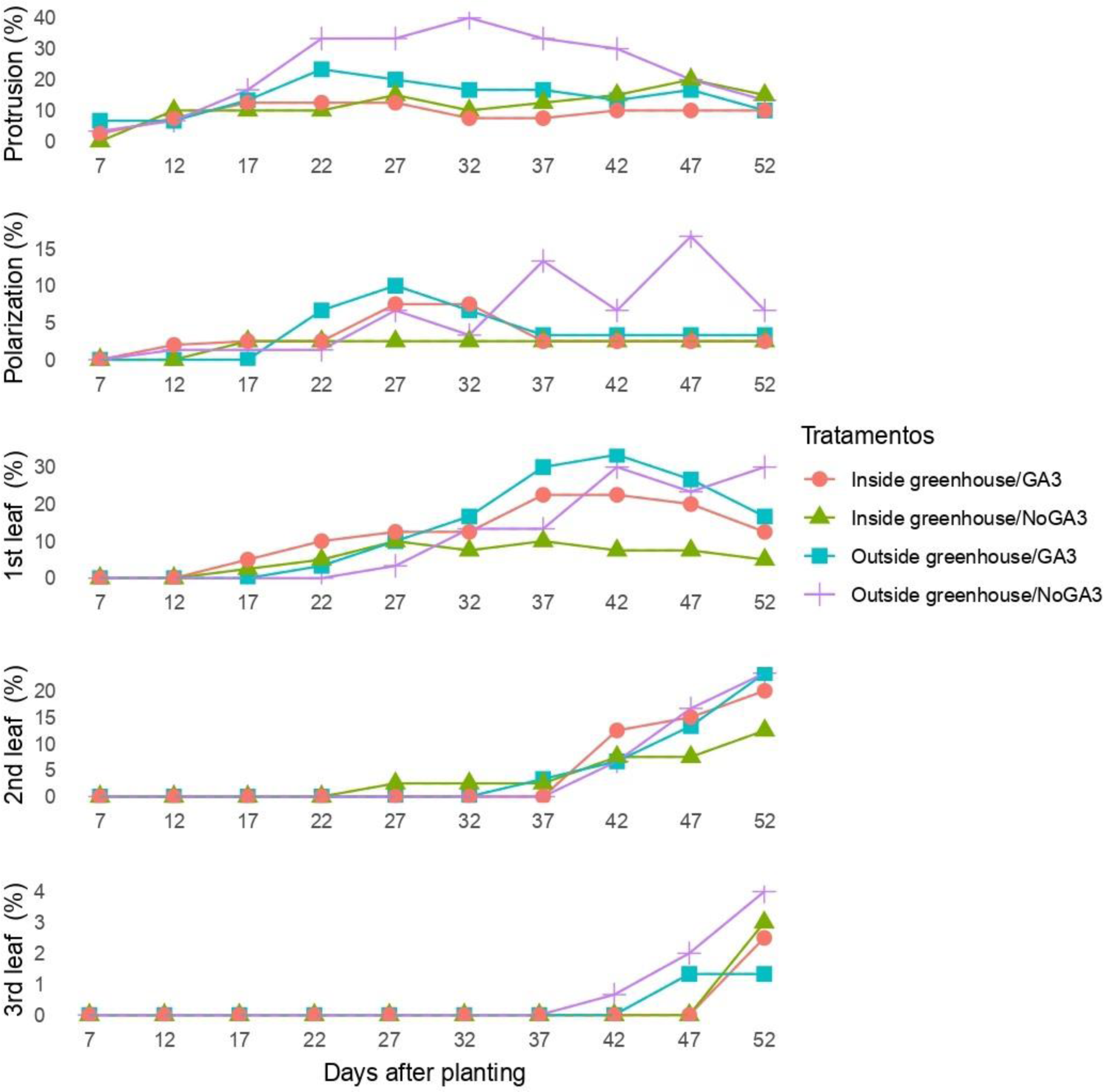
Stages of germination and post-seminal development of the “sempre-viva” *Janeirona carrasqueira* (morphotype of *Comanthera bisulcata*). Seeds collected in the first week of February 2020 in an open seedbed (outside the greenhouse) and inside the greenhouse, treated or not with GA3.

Considering the values of relative frequency of germination, it can be seen that regardless of where the seeds were collected, germination was distributed over time, with several germination peaks, presenting a polymodal behavior. However, the first and highest germination peaks were observed on the 22^nd^ day after sowing for the seeds collected in open-air beds, and on the 12^th^ day after sowing for seeds collected inside the greenhouse. Germination stabilization occurred approximately 47 d after sowing (Fig. 5).

**Figure 5.**
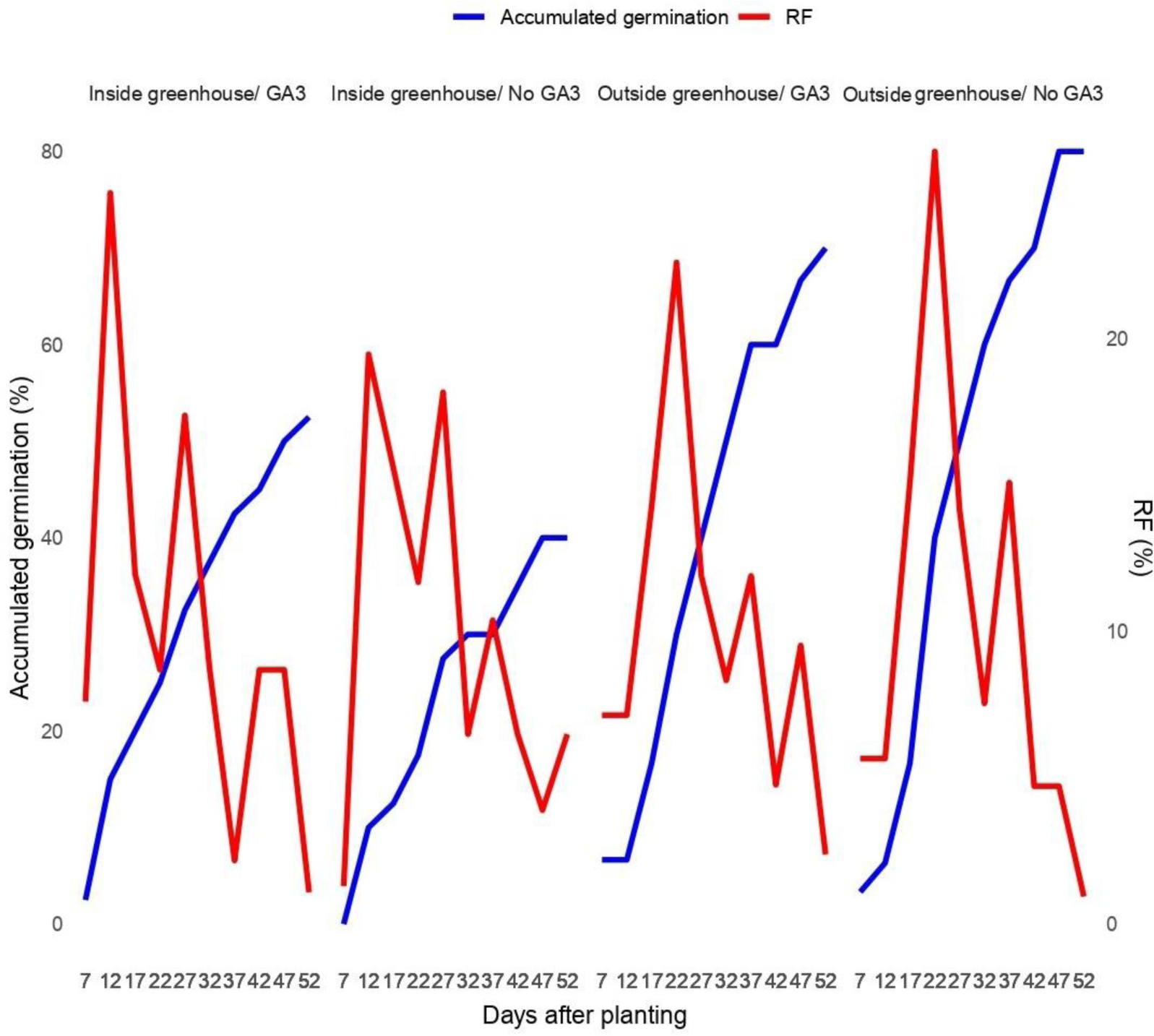
Accumulated germination rate and relative frequency of seed germination of the “sempre-viva” *Janeirona carrasqueira* (morphotype of *Comanthera bisulcata*), in seeds collected in the first week of February 2020, in plants grown in open-air beds and in greenhouses. Nt = Total number of germinated seeds.

Germination rates were significantly higher (p<0.05) in the treatments using seeds collected outside the greenhouse (natural pollination treatment), with a median of 80% for seeds without GA3 application and 70% for seeds immersed in GA3 (Fig. 6). The germination rate of seeds resulting from natural pollination was 100% higher than that of seeds produced by plants grown in a greenhouse (autogamy).

**Figure 6.**
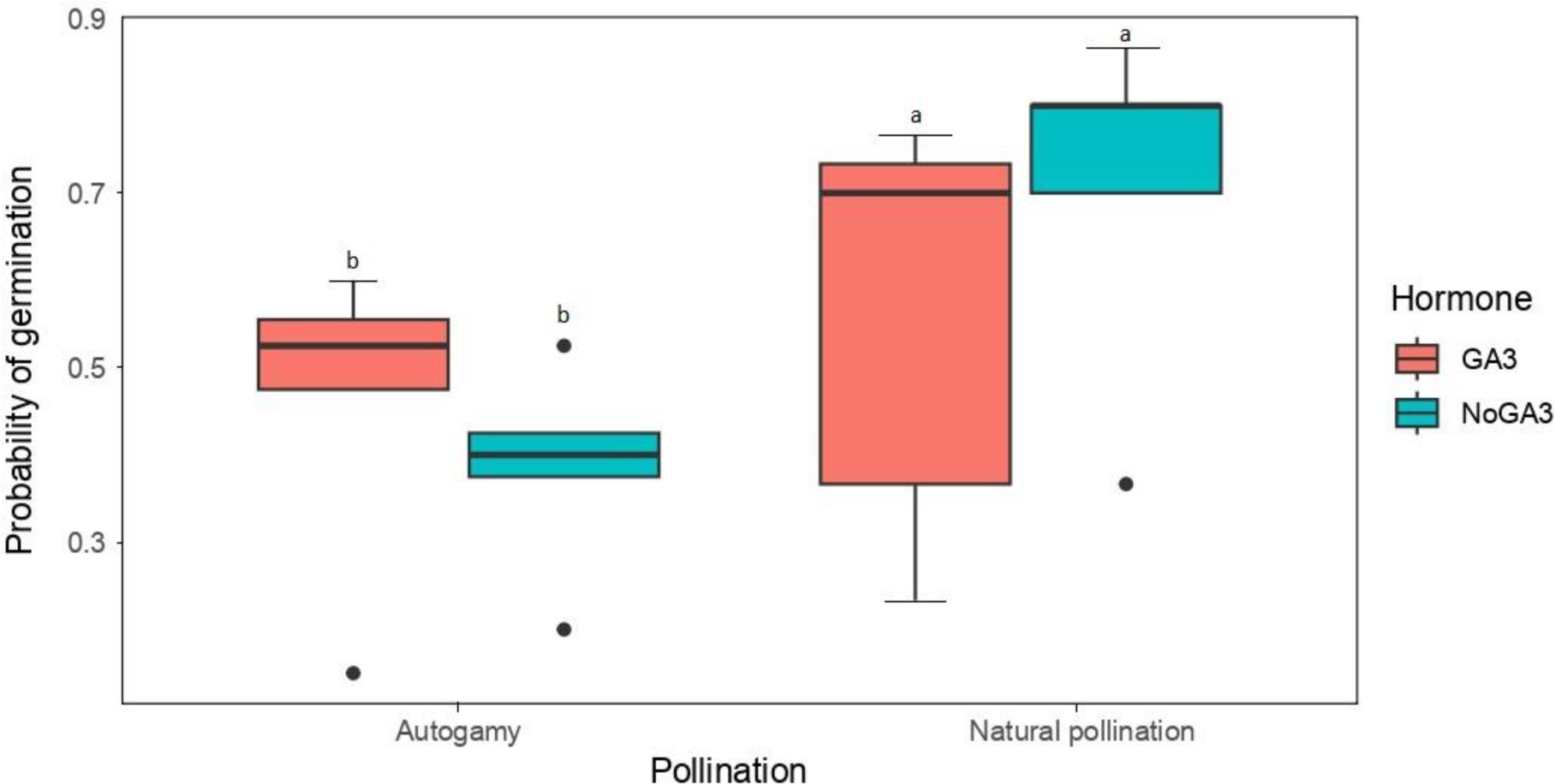
Distribution of the germination rate of Janeirona carrasqueira (morphotype of *Comanthera bisulcata*), in seeds collected outside and inside the greenhouse and treated with GA3 or without GA3. Vertical lines denote minimum and maximum values of germination rates; gray bars span the first and third quartiles divided by the median. The points are outliers.

The model that best explains the results of the germination test was the one that included only the pollination mode as a predictor variable, with the lowest dAIC value (0.0) and the highest weight (0.485) (Table 1). The complete models, both with interaction and without interaction, had similar AIC values. This indicates that the inclusion of interaction or its exclusion did not result in a substantial difference in the model’s goodness of fit. On the other hand, the null model and the model with only the hormone had the worst performance, suggesting that the effect of the hormone on the germination rate is weak. Thus, the simple model, considering only pollination, was more adequate to explain the observed results.

**Table 1.** Results of model selection analysis for germination succes in seeds of *Janeirona carrasqueira* (morphotype of *Comanthera bisulcata*) subjected to two pollination treatments (natural x autogamy) and hormone induction (with and without giberelin application).

| Model | Germination Success |  |  |
| --- | --- | --- | --- |
|  | dAIC | DF | Weight |
| **Complete model with interaction** | 1.7 | 5 | 0.212 |
| **Complete model without interaction** | 1.7 | 4 | 0.203 |
| **Pollination Mode only** | 0.0 | 3 | 0.485 |
| **Hormone only** | 5.6 | 3 | 0.029 |
| **Null Model** | 3.8 | 2 | 0.071 |

The coefficients of fixed effects of the selected optimal model (Table 2) show the influence of the predictor variables on the response variable. The estimated coefficient for the variable “Natural Pollination” was positive (0.9620) and statistically significant (Pr(>|z|) = 0.00998*), indicating a favorable impact on the germination rate. In other words, natural pollination resulted in a higher proportion of germinated seeds compared to autogamy. The average probability of germination estimated by the model for the plants located inside the greenhouse (Pollination = 0, ie, Autogamy) was approximately 41.33%. Enquanto que para as plantas sujeitas à polinização natural, a probabilidade média estimada de germinação foi de aproximadamente 65% (Fig. 7).

**Figure 7.**
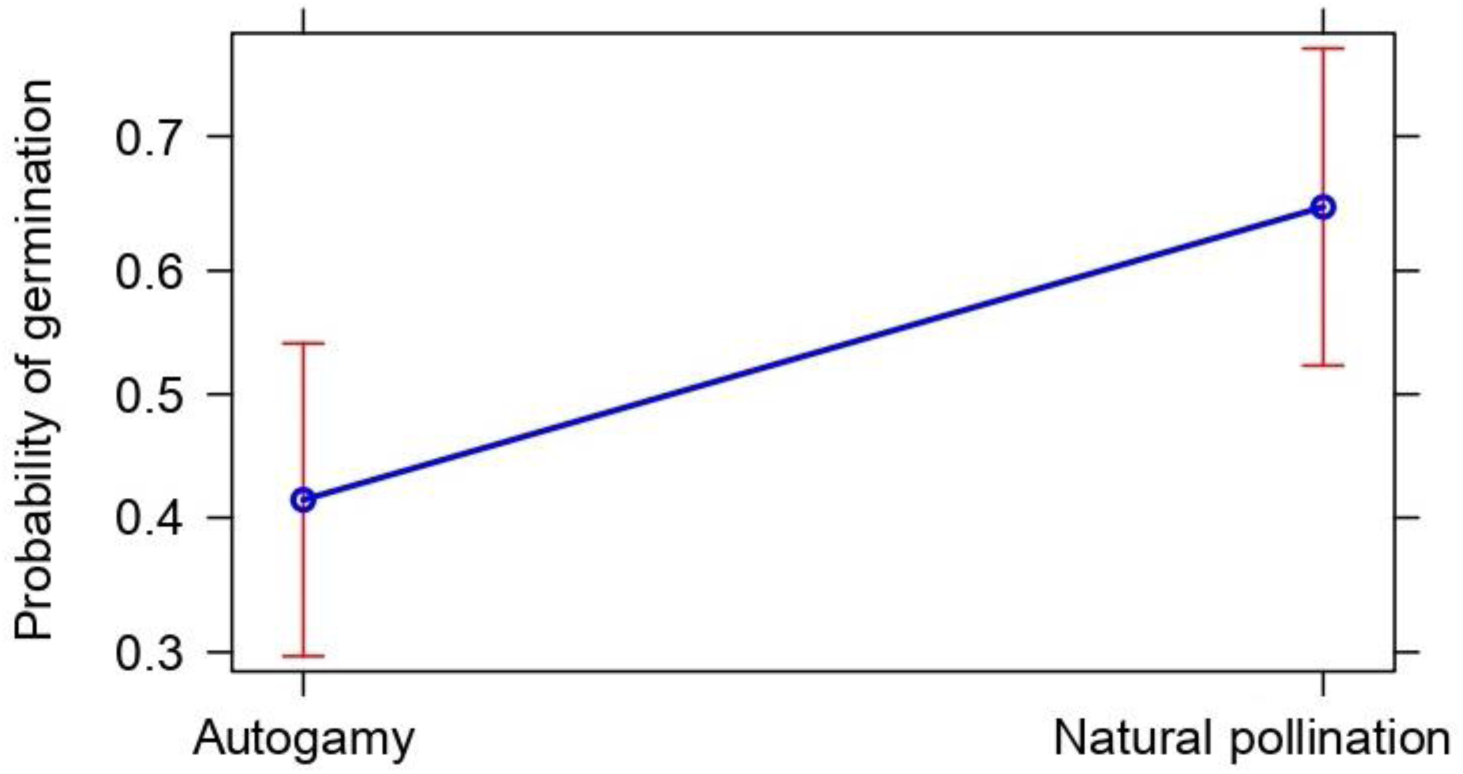
Generalized Mixed Model (GLMM) prediction for the effect of pollination on the germination of Janeirona carrasqueira (morphotype of *Comanthera bisulcata*).

**Table 2.** Summary of the model selected to estimate germination success in seeds of Janeirona carrasqueira (morphotype of *Comanthera bisulcata*), with emphasis on the coefficients of Fixed and Random Effects.

| Fixed effects |  |  |  |  |
| --- | --- | --- | --- | --- |
|  | Estimate | Std. Error | z value | Pr(>\|z\|) |
| **Intercept** | -0.3471 | 0.2620 | -1.325 | 0.18531 |
| **PollinationNatural pollination** | 0.9620 | 0.3734 | 2.576 | 0.00998* |
| Random effects |  |  |  |  |
| Groups | Name | Variance | Std. Dev. |  |
| **Rep** | Intercept | 0.5356 | 0.7318 |  |

## 4. Discussion

Our study showed that the *Comanthera bisulcata* morphotype is self-compatible and present a generalized insect pollination system (even though some insects do not carry pollen) with pollen and nectar as rewards. We also demonstrated a relationship between the type of pollination and the physiological mechanisms of germination, where the most influential factor was the pollination system, not the application of the inducer nor the interaction of pollination system and hormone application.

### 4.1. Floral visitors and pollinators

The “sempre-viva” *Janeirona carrasqueira* produces nectar and pollen as floral resources and attracts a wide range of small insects as floral visitors. Other *Comanthera* species studied in Campo Rupestre (*C. curralensis*, *C. mucugensis, C. elegans*) also exhibited pollination by small insects of the orders Hymenoptera, Diptera, and Coleoptera, reaffirming the exitence of a generalist pollination systems in the genus (Ramos et al., 2005; Oriani et al., 2009). In *C. elegans*, pollinator visitation peaks coincided with periods of high temperature and low relative humidity, corresponding to complete opening of the capitula (Oriani et al., 2009). This occurs because of the movement of the involucral bracts, which are influenced by humidity (Oriani and Scatena, 2009).

Nectar availability at the beginning of anthesis may be a pollen-saving mechanism. Thus, the consumption pressure on this resource is reduced, resulting in a lower probability of visits by insects that collect pollen at the beginning of anthesis (Fig. 2A). This strategy may also maximize the chances of transport between flowers and, consequently, pollination, as visits by insects that collect pollen are more likely when more flowers are visited. Furthermore, dispersing the pollen with nectar-eating visitors increases the chance that it will be transferred to another flower and will not become food for adult flies or bee larvae (the main pollen collectors). This becomes especially important because the pollen stock within a flower is limited, and the nectar supply can be renewed through the production of this resource throughout the day (Brito et al., 2017). In addition, female flowers exclusively offer nectar, which further increases the potential for floral visitors who exclusively collect pollen not to act as pollinators because they would have no reason to go to the female flowers.

### Germination test

The germination rate of this species was greater than 40% in both the treatments. However, *Janeirona carrasqueira* seeds collected outside the greenhouse (natural pollination treatment) attained the highest germination rates (above 70%). This demonstrates that in this environment, the seeds did not show any type of primary dormancy. This also indicated that natural pollination contributed more effectively to the reproductive success of the species (Ramos et al., 2005; Oriani et al., 2009; Del-Claro et al., 2019; Martins Junior et al., 2022).

The small size of the seeds of the genus *Comanthera* allows for an increase in the number of seeds produced and their dispersion capacity. This, in turn, increases the chance of finding favorable locations for species establishment (Echternacht et al., 2014). This species first developed leaves and then roots, which classified the seeds as positive photoblastic. Positive photoblastism is an important strategy that prevents the germination of small buried seeds because the energy reserves would be insufficient to reach the surface (Garcia et al., 2020). Leaf development, one of the first post-germination stages, allows the species to obtain the energy necessary for its establishment through photosynthesis (Mascarenhas and Scatena, 2021). Although small, the seed reserve of the *C. bisulcata* morphotype was sufficient for the emission of three leaves at 42 d after sowing.

Species germination was distributed over time for both treatments. Polymodal germination behavior can be attributed to the fact that seeds of native species often exhibit high genetic variability, which represents a crucial adaptive strategy for their survival (Gonçalves-Magalhães et al., 2021). Thus, the seeds that did not germinate at the end of the period with favorable conditions reduce their metabolism, activating it in adequate environmental conditions for their establishment (Gonçalves-Magalhães et al., 2021). This indicates that this species can establish seed banks in the soil, which occur in most species from Campo Rupestre (Garcia et al., 2014; Garcia et al., 2020).

Autogamy was demonstrated by the formation of seeds in the capitulum inside the greenhouse. That may be a result of self-pollination, a common phenomena in Eriocaulaceae happening due to the spatial proximity and temporal overlap of male and female flower maturation in inflorescences (Oriani et al., 2009; Horiuchi et al., 2020). Species that spontaneously self-pollinate and are self-compatible can produce seeds without the need for a pollination vector, which is advantageous for adapting to certain environments, especially degraded ones (Proctor et al., 1996). Thus, mixed mating strategies allow reproduction by crossing when pollinators are abundant or by geitonogamy/apomixis when pollinators are scarce or absent (Martins Junior et al., 2022). Thus, this species can reproduce even in years that are unfavorable to pollinators. However, as cross-pollination brings a lot of genetic diversity and the species lives in reasonably inhospitable environments, there is a clear difference between the germination and vigor of seeds produced by cross-pollination and autogamy.

In *Comanthera elegans*, a lower percentage of germination was observed for seeds resulting from autogamy, which may be considered a consequence of inbreeding depression (Oriani et al., 2009; Horiuchi et al., 2021). According to the authors, this could be explained by the harmful effects of homozygous genes resulting from inbreeding. Therefore, entomophily is the most efficient way for this species to reproduce. Ramos et al. (2005) also considered entomophily as the main mode of pollination in *Comanthera mucugensis* and *Comanthera curralensis* based on research on floral visitors and insect activities in flowers, although autogamy also occurred in these species. Mixed breeding systems have been also reported for *Paepalanthus* sps.(*P. bifidus* and *P. tortilis*) (Martins Junior et al., 2022).

The low germination rate of autogamy seeds may be related to dormancy. Seed germination and dormancy are physiological processes that are regulated by endogenous hormones, of which abscisic acid (ABA) and gibberellins (GA) stand out (Garcia et al., 2020). ABA is a positive regulator of dormancy, whereas gibberellins are involved in breaking dormancy and promoting germination (Barreto et al., 2020). Thus, treatments that alter the hormonal balance by decreasing ABA levels, increasing GA levels, or both contribute to overcoming physiological dormancy (Baskin and Baskin, 2004).

However, despite gibberellins being promoters of dormancy breakage and germination induction, for “*Janeirona carrasqueira*” seeds, the use of GA3 was not significantin in germination. Thus, we suggest that this species may be less sensitive to the exogenous application of gibberellins. In *Syngonanthus verticillatus*, the use of gibberellin was not efficient for germination; in contrast, seeds treated with fluridone, an ABA synthesis inhibitor, were effective in overcoming dormancy (Barreto et al., 2020). The low responsiveness to gibberellin can also be indicative of another type of numbness that is not physiological but morphological.

Pollinator-friendly management practices are essential for ensuring effective cross-pollination during the *in situ* and *ex situ* conservation of Eriocaulaceae (Horiuchi et al., 2021). Studies reporting the occurrence of entomophily in everlasting flower species are very important for conservation management, given the practice of recurrent burning in areas where these species occur (Ramos et al., 2005). According to Ramos et al. (2005), this practice affects the reproduction of these species, as many of their floral visitors are resident species with short flight characteristics that can lead to a decrease in the number of potential pollinators. However, it should be considered that the controlled burning used in the management of everlasting flowers species, besides being restricted to the locations where collected species occur, is not conducted during the period when the plants are in the pollination phase. Therefore, certain associations between fire management and its effects on pollinators must be established with caution. Finnaly, the data from this study present relevant contributions to the understanding of the biology of *Comanthera bisulcata*, playing a crucial role in the formulation of management strategies aimed at the effective conservation of the species.

## 5. Conclusion

For the first time, we demonstrate that the pollination system can play a decisive role for Eriocaulaceae, both in seedling germination and vigor, with effects beyond physiology. Considering the importance of the reproductive system in germination, together with the generalist pollination system and the dynamics of seedling birth, we emphasize the need to incorporate plant reproduction in management plans for traditional flower picking. This action is fundamental both in the implementation of appropriate measures for *in situ* conservation, and in the cultivation of evergreens (which may contribute to reducing collection pressure in areas where the species occurs naturally).

## Acknowledgements

This work was carried out with the support of the Coordination for the Improvement of Higher Education Personnel (CAPES) - Financing Code 001 and CNPq (Proc. 400904/2019-5). ARR, MLA and JCB are also grateful to CNPq (Proc. 423939/2021-1, Proc. 311665/2022-5, Proc. 441984/2018-5 and Proc. 400904/2019-5) and FAPEMIG (APQ 0932-21).

## References

Abrahão, A., Costa, PDB., Lambers, H., Andrade, SAL., Sawaya, ACHF., Ryan, MH., Oliveira, RS, 2019. Soil types select for plants with matching nutrient-acquisition and-use traits in hyperdiverse and severely nutrient-impoverished campos rupestres and cerrado in Central Brazil. Journal of Ecology, 107, 1302–1316. https://doi.org/10.1111/1365-2745.13111.

Alvares, CA., Stape, JL., Sentelhas, PC., Gonçalves, JDM., Sparovek, G., 2013. Köppen’s climate classification map for Brazil. Meteorologische Zeitschrift, 22, 711–728. https://doi.org/10.1127/0941-2948/2013/0507.

Andrino, CO., Costa, FN., Sano, PT., 2015. O gênero Paepalanthus Mart. (Eriocaulaceae) no Parque Estadual do Biribiri, Diamantina, Minas Gerais, Brasil. Rodriguésia, 66, 393–419. https://doi.org/10.1590/2175-7860201566209.

Andrino, CO., Santana, PC., Lovo, J., Barbosa-Silva, RG., Albuquerque-Lima, S., Zappi, DC., 2022. Anthers in blue: A hidden rhapsody in Amazonian Eriocaulaceae. Ecology, 103, e3636. https://doi.org/10.1002/ecy.3636.

Barreto, LC., Herken, D., Silva, BM., Munné-Bosch, S., Garcia, QS., 2020. ABA and GA 4 dynamic modulates secondary dormancy and germination in Syngonanthus verticillatus seeds. Planta, 251, 1–10. https://doi.org/10.1007/s00425-020-03378-2.

Baskin, JM., Baskin, CC., 2004. A classification system for seed dormancy. Seed Science Research, 14, 1–16. https://doi.org/10.1079/SSR2003150.

Borges, T., Branford, S, 2021. Brazil: the flowers of sustainability. Latin America Bureal. https://lab.org.uk/brazil-the-flowers-of-sustainability/ (accessed on 4 July 2023).

Brito, VL., Rech, AR., Ollerton, J., Sazima, M., 2017. Nectar production, reproductive success and the evolution of generalised pollination within a specialised pollen-rewarding plant family: a case study using Miconia theizans. Plant Systematics and Evolution, 303, 709–718. https://doi.org/10.1007/s00606-017-1405-z.

Castro, SA., Silveira, FA., Marcato, MS., Lemos-Filho, JP., 2018. So close, yet so different: Divergences in resource use may help stabilize coexistence of phylogenetically-related species in a megadiverse grassland. Flora, 238, 72–78. https://doi.org/10.1016/j.flora.2016.11.018.

Colli-Silva, M., Vasconcelos, TNC., Pirani, J., 2019. Outstanding plant endemism levels strongly support the recognition of campo rupestre provinces in mountaintops of eastern South America. Journal of Biogeography, 46, 1723–1733. https://doi.org/10.1111/jbi.13585.

Costa, FN., Trovó, M., Sano, PT., 2008. Eriocaulaceae na Cadeia do Espinhaço: riqueza, endemismo e ameaças. Megadiversidade, 4, 117–125.

Del-Claro, K., Rodriguez-Morales, D., Calixto, ES., Martins, AS., Torezan-Silingardi, H.M., 2019. Ant pollination of Paepalanthus lundii (Eriocaulaceae) in Brazilian savanna. Annals of Botany, 123, 1159–1165. https://doi.org/10.1093/aob/mcz021.

Echternacht, L., Sano, PT., Bonillo, C., Cruaud, C., Couloux, A., Dubuisson, JY., 2014. Phylogeny and taxonomy of Syngonanthus and Comanthera (Eriocaulaceae): Evidence from expanded sampling. Taxon, 63, 47–63. https://doi.org/10.12705/631.36.

Echternacht, L., Trovó, M., Costa, F.N., Sano, P.T., 2012. Análise comparativa da riqueza de Eriocaulaceae nos parques estaduais de Minas Gerais, Brasil. MGBiota, 4, 18–31.

FAO (Food and agriculture organization of the United Nations), 2020. Apanhadoras e apanhadores de flores sempre-vivas recebem reconhecimento internacional da FAO como o primeiro Patrimônio Agrícola Mundial do Brasil. http://www.fao.org/brasil/noticias/detail-events/en/c/1265788/ (accessed 15 April 2023).

Fernandes, GW., Arantes-Garcia, L., Barbosa, M., Barbosa, NP., Batista, EK., Beiroz, W., … Silveira, FA., 2020. Biodiversity and ecosystem services in the Campo Rupestre: A road map for the sustainability of the hottest Brazilian biodiversity hotspot. Perspectives in Ecology and Conservation, 18, 213–222. https://doi.org/10.1016/j.pecon.2020.10.004.

Garcia, QS., Barreto, LC., Bicalho, EM., 2020. Environmental factors driving seed dormancy and germination in tropical ecosystems: A perspective from campo rupestre species. Environmental and Experimental Botany, 178, 104164. https://doi.org/10.1016/j.envexpbot.2020.104164.

Garcia, QS., Oliveira, PG., Duarte, DM., 2014. Seasonal changes in germination and dormancy of buried seeds of endemic Brazilian Eriocaulaceae. Seed Science Research, 24, 113–117. https://doi.org/10.1017/S0960258514000038.

Giulietti, AM., Andrade, MJG., Scatena, VL., Trovó, M., Coan, AI., Sano, PT., Van den Berg, C., 2012. Molecular phylogeny, morphology and their implications for the taxonomy of Eriocaulaceae. Rodriguésia, 63, 01–19. https://doi.org/10.1590/S2175-78602012000100001.

Gonçalves-Magalhães, C., De Santana, DG.; Ribeiro-Oliveira, JP., 2021. Misunderstanding on germination sensu stricto leads us to a false positive germination-dormancy balance in diaspores of Paepalanthus chiquitensis Herzog (Eriocaulaceae), a threatened everlasting flowering species. Plant Species Biology, 36, 246–257. https://doi.org/10.1111/1442-1984.12319.

Horiuchi, Y., Kamijo, T., Tanaka, N., 2020. Biological and ecological constraints to the reintroduction of Eriocaulon heleocharioides (Eriocaulaceae): A species extinct in the wild. Journal for Nature Conservation, 56, 125866. https://doi.org/10.1016/j.jnc.2020.125866.

Horiuchi, Y., Kamijo, T., Tanaka, N., 2021. Floral and pollination characteristics of Eriocaulon heleocharioides, an extinct species in the wild, for evidence-based conservation management. Plant Biology, 23, 546–555. https://doi.org/10.1111/plb.13236.

INMET (Instituto Nacional de Meteorologia), 2020. Normais Climatológicas do Brasil 1991-2020. https://portal.inmet.gov.br/normais (Accessed 20 March 2023).

Labouriau, LG., Valadares, MEB., 1976. On the germination of seeds Calotropis procera (Ait.) Ait.f. Anais da Academia Brasileira de Ciências, Rio de Janeiro. 48, 263–284.

Martins Junior, ER., da Costa, ACG., Milet-Pinheiro, P., Navarro, D., Thomas, WW., Giulietti, AM., Machado, IC., 2022. Mixed pollination system and floral signals of Paepalanthus (Eriocaulaceae): insects and geitonogamy ensure high reproductive success. Annals of Botany, 129, 473–484. https://doi.org/10.1093/aob/mcac008.

Mascarenhas, AAS., Scatena, VL., 2021. Seed morphology and post-seminal development in Leiothrix Ruhland (Eriocaulaceae, Poales). Feddes Repertorium, 132, 9–19. https://doi.org/10.1002/fedr.202000022.

MMA (Ministério do Meio Ambiente), 2022. Lista Nacional de Espécies Ameaçadas de Extinção (Portaria MMA N°. 148, de 07 de Junho de 2022). https://www.in.gov.br/en/web/dou/-/portaria-mma-n-148-de-7-de-junho-de-2022-406272733. (Accessed 20 June 2023).

Oliveira, MNS., Cruz, LI., Tanaka, MK., 2014. Collection time and seed germination of commercialized Comanthera (Eriocaulaceae) from Serra do Ambrósio, Minas Gerais. Brazilian Journal of Botany, 37, 19–27. https://doi.org/10.1007/s40415-013-0045-y.

Oliveira, PG., Garcia, QS., 2011. Germination characteristics of Syngonanthus seeds (Eriocaulaceae) in campos rupestres vegetation in south-eastern Brazil. Seed Science Research, 21, 39–45. https://doi.org/10.1017/S0960258510000346.

Oriani, A., Sano, PT., Scatena, VL., 2009. Pollination biology of Syngonanthus elegans (Eriocaulaceae-Poales). Australian Journal of botany, 57, 94–105. https://doi.org/10.1071/BT08119.

Oriani, A., Scatena, VL., 2009. The movement of involucral bracts of Syngonanthus elegans (Eriocaulaceae-Poales): anatomical and ecological aspects. Flora-Morphology, Distribution, Functional Ecology of Plants, 204, 518–527. https://doi.org/10.1016/j.flora.2008.07.003.

Parra, LR., Giulietti, AM., Andrade, MJG,. Berg, C., 2010. Reestablishment and a new circumscription of Comanthera (Eriocaulaceae). Taxon, 59, 1135–1146. https://doi.org/10.1002/tax.594013.

Pawłowski, TA., 2009. Proteome analysis of Norway maple (Acer platanoides L.) seeds dormancy breaking and germination: influence of abscisic and gibberellic acids. BMC Plant Biol. 9, 48–61. https://doi.org/10.1186/1471-2229-9-48.

Proctor, M., Yeo, P., Lack, A., 1996. The natural history of pollination. Harper Collins, London. 479p.

R Core Team., 2013. R: A language and environment for statistical computing. R Foundation for Statistical Computing.

Ramos, COC., Borba, EL., Funch, LS., 2005. Pollination in Brazilian Syngonanthus (Eriocaulaceae) species: evidence for entomophily instead of anemophily. Annals of botany, 96, 387–397. https://doi.org/10.1093/aob/mci191.

Schmidt, IB., Figueiredo, IB., Borghetti, F., Scariot, A., 2008. Produção e germinação de sementes de “capim dourado”, Syngonanthus nitens (Bong.) Ruhland (Eriocaulaceae): implicações para o manejo. Acta botanica brasilica, 22, 37–42. https://doi.org/10.1590/S0102-33062008000100005.

Silveira, FA., Negreiros, D., Barbosa, NP., Buisson, E., Carmo, FF., Carstensen, DW., … Lambers, H., 2016. Ecology and evolution of plant diversity in the endangered campo rupestre: a neglected conservation priority. Plant and soil, 403, 129–152. https://doi.org/10.1007/s11104-015-2637-8.

Simões, FC., Oliveira Paiva, PD., Tavares, TS., Paiva, R., 2007. Germinação de sementes de sempre-vivas (Syngonanthus elegans e S. venustus). Revista Brasileira de Horticultura Ornamental, 13, 79–83. https://doi.org/10.14295/rbho.v13i1.207.

Stützel, T., Trovó, M., 2013. Inflorescences in Eriocaulaceae: taxonomic relevance and practical implications. Annals of Botany, 112, 1505–1522. https://doi.org/10.1093/aob/mct234.

Sun, TP., Gubler, F., 2004. Molecular mechanism of gibberellin signaling in plants. Annual Review of Plant Biology, 55, 197–223. https://doi.org/10.1146/annurev.arplant.55.031903.141753.

Tanaka, N., Ono, H., Nagata, S., 2015. Floral visitors of Eriocaulon heleocharioides (Eriocaulaceae), an extinct aquatic species in the wild. Bulletin of the National Museum of Natural Sciences, Series B, 41, 179–182.

Vasconcelos, TN., Alcantara, S., Andrino, CO., Forest, F., Reginato, M., Simon, MF., Pirani, JR., 2020. Fast diversification through a mosaic of evolutionary histories characterizes the endemic flora of ancient Neotropical mountains. Proceedings of the Royal Society B, 287, 20192933. https://doi.org/10.1098/rspb.2019.2933.

Vieira, JP., de Souza, MJ., Teixeira, JM., Carvalho, FPD., 2010. Estudo da precipitação mensal durante a estação chuvosa em Diamantina, Minas Gerais. Revista Brasileira de Engenharia Agrícola e Ambiental, Campina Grande, 14, 762–767. https://doi.org/10.1590/S1415-43662010000700012.

